# Nutrigonometry I: using right-angle triangles to quantify nutritional trade-offs in multidimensional performance landscapes

**DOI:** 10.1101/2021.11.25.469978

**Authors:** Juliano Morimoto, Pedro Conceição, Christen Mirth, Mathieu Lihoreau

**Affiliations:** School of Biological Sciences, University of Aberdeen, Zoology Building, Tillydrone Ave, Aberdeen AB24 2TZ; Institute of Mathematics, University of Aberdeen, King’s College, Aberdeen AB24 3FX; School of Biological Sciences, Monash, University, Melbourne, Victoria, Australia; Research Center on Animal Cognition (CRCA), Center for Integrative Biology (CBI); CNRS, University Paul Sabatier – Toulouse III, France

**Author notes:** Correspondence: Dr Juliano Morimoto. **Authors’ contributions** JM conceptualised the nutrigronometry model, coded the scripts to conduct the analysis, and wrote the first draft of the manuscript. JM and PC implemented the use of persistence homology. All authors contributed to the revising and editing of the manuscript and approved its submission.

**Keywords:** Nutritional Geometry, trigonometry, lifespan-reproduction trade-off, fitness maps, *Drosophila melanogaster*

## Abstract

Animals regulate their diet in order to maximise the expression of fitness traits that often have different nutritional needs. These nutritional trade-offs have been experimentally uncovered using the Geometric framework for nutrition (GF). However, current analytical methods to measure such responses rely on either visual inspection or complex models applied to multidimensional performance landscapes, making these approaches subjective, or conceptually difficult, computationally expensive, and in some cases inaccurate. This limits our ability to understand how animal nutrition evolved to support life-histories within and between species. Here, we introduce a simple trigonometric model to measure nutritional trade-offs in multidimensional landscapes (‘Nutrigonometry’). Nutrigonometry is both conceptually and computationally easier than current approaches, as it harnesses the trigonometric relationships of right-angle triangles instead of vector calculations. Using landmark GF datasets, we first show how polynomial (Bayesian) regressions can be used for precise and accurate predictions of peaks and valleys in performance landscapes, irrespective of the underlying structure of the data (i.e., individual food intakes *vs* fixed diet ratios). Using trigonometric relationships, we then identified the known nutritional trade-off between lifespan and reproductive rate both in terms of nutrient balance and concentration. Nutrigonometry enables a fast, reliable and reproducible quantification of nutritional trade-offs in multidimensional performance landscapes, thereby broadening the potential for future developments in comparative research on the evolution of animal nutrition.

## Introduction

Animals often require different nutrient blends to maximize concurrent life-history traits, creating the potential for a conflict between nutrient allocation (Simpson and Raubenheimer 2012; Raubenheimer and Simpson 2020). When the optimum nutrition for several traits cannot be achieved simultaneously, a compromise in feeding decisions must exist in order to support the expression of one trait over another (‘nutritional trade-off’) (Lee et al. 2008; Maklakov et al. 2008). Previous research has identified nutritional trade-offs between lifespan and reproduction or between immunity and reproduction across many different taxa including *D. melanogaster* (Lee et al. 2008; Ponton et al. 2015), tephritid fruit flies (Fanson and Taylor 2012; Fanson et al. 2012), crickets (Maklakov et al. 2008; Rapkin et al. 2018; Guo et al. 2021; Treidel et al. 2021) and mice (Solon-Biet et al. 2014) [see also reviews by (Ponton et al. 2011; Schwenke et al. 2016)]. Even traits related to different aspects of the same life-history can vary in nutritional requirements during the lifetime of an animal, as seen for pre- and post-mating traits related to reproduction of many insect species such as sperm number and viability (Bunning et al. 2015), fertilization success across sperm competitive contexts (Morimoto and Wigby 2016), cuticular hydrocarbons, courtship song and sperm viability (Ng et al. 2018) as well as size and numbers of eupyrene and apyrene sperms (Gage and Cook 1994). Thus, nutritional trade-offs are likely ubiquitous and impose significant constraints on the feeding behaviour of individuals.

Measuring nutritional trade-offs can be challenging because of the interactive effects of nutrient ratios and concentrations on the expression of life-histories (Stearns 1992; Roff 2002; Hunt et al. 2004; Simpson and Raubenheimer 2012). In the last decades, however, a method known as Geometric Framework of Nutrition (GF) has emerged as a powerful unifying framework capable of disentangling the multidimensional effects of nutrients (both ratios and concentrations) on life-history traits and fitness (Simpson and Raubenheimer 1993*a*). The GF has been applied to a diverse range of nutritional studies across species such as flies (Lee et al. 2008; Reddiex et al. 2013; Jensen et al. 2015; Ponton et al. 2015; Morimoto and Wigby 2016; Kutz et al. 2019) (Fanson and Taylor 2012; Fanson et al. 2012) (Barragan-Fonseca et al. 2018, 2021), crickets (Ng et al. 2018) (Rapkin et al. 2018) (Maklakov et al. 2008), cockroaches (Bunning et al. 2015), domestic cats and dogs (Hewson-Hughes et al. 2011, 2013), and mice (Solon-Biet et al. 2014; Morimoto et al. 2019), being paramount for advancing our understanding of complex physiological and behavioural processes across ecological environments and even human health (Simpson et al. 2017). As a result, developing a simple, intuitive, and accurate quantitative method for quantifying nutritional trade-offs has become a key issue for comparative nutrition, which will allow new avenues of research for insights into the evolution of physiological and behavioural modulation of nutritional responses (Morimoto and Lihoreau 2020).

Recent initiatives have been made but these are complex to navigate and therefore studies continue to be published with visual inspection or with inaccurate methods to quantify the strength of nutritional trade-offs in GF landscapes [e.g., (Polak et al. 2017; Ng et al. 2018, 2019; Kutz et al. 2019; Ma et al. 2020; Barragan-Fonseca et al. 2021)]. Why is it so difficult to measure nutritional trade-offs in GF multidimensional fitness landscapes? The fundamental limitation in all models so far is identifying and delimitating the region of interest (i.e., peaks and, to a lesser extent, valleys) for comparisons of distances between peaks of different traits (or same trait between species). For instance, (Rapkin et al. 2018) proposed the use of regression slopes (rather than the coordinates of the optimum) nutrients onto the fitness trait *i* as coordinates of a vector 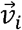. From this, the angle *θ*′ between vectors 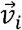 and 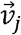 for traits *i* and *j*, respectively, can be calculated as the estimate of the strength of the nutritional trade-off. However, in this approach, the domain of each vector coordinate is all real numbers ℝ even though the domain of the fitness landscape is constrained to all positive real numbers ℝ^+^. Consequently, this violates the domain constraints of the nutritional space upon which GF is performed, which in turn result in overestimation of the strength of nutritional trade-offs (Morimoto and Lihoreau 2019). To address this limitation, we proposed to use the coordinates of the peak (or valley) from the nutritional space as vector coordinates for the position vector 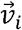, from which the angle *θ* between position vectors 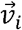 and 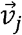 for traits *i* and *j*, respectively, can be estimated as measure of the strength of the nutritional trade-off. This overcomes the violation of domains between vector coordinates and the nutritional space [in (Rapkin et al. 2018)] and therefore ensures that the estimate of *θ* is calculated in the same domain of the GF fitness landscapes. However, this Vector of Positions approach relies on the peak estimates from a SVM machine learning model which is computationally expensive particularly in *n* dimensions, where n > 3, and sensitive to the characteristics of the input data (e.g., if the data contains food intakes as in (Lee et al. 2008; House et al. 2015; Jensen et al. 2015; Morimoto and Lihoreau 2019) or a grid of fixed diet ratios as in (Kutz et al. 2019)], identifying local as well as global peaks that introduce noise into the analysis (shown here). Albeit useful, the Vector of Positions approach can be cumbersome to implement across different datasets, computationally expensive to obtain estimates of peaks (or valleys), and as a result can generate inaccurate estimates of the strength of nutritional trade-offs. Thus, to date, there are no proposed solutions that address the above limitations, which creates a significant bottleneck in studies of nutrition that limits the multidimensional power of the GF framework.

Here, we address the limitations of current models by proposing a simpler framework (Nutrigonometry) upon which the strength of nutritional trade-offs can be calculated in 3D fitness landscapes, irrespective of the structure of the nutritional data to analyze. Using landmark GF datasets, we first investigated the performance of different ‘off-the-shelf’ machine learning models in predicting the peak in the fitness landscapes, in order to find the most accurate and computationally inexpensive model. We achieved this by integrating several measurements of predictive error, variance in predicted peak area, and topological characteristics of the predicted peak region. Next, we used simple trigonometric functions and relationships to estimate the strength of nutritional trade-offs both in terms of nutrient balance as well as nutrient concentration, or both. Our approach opens new avenues of research in multidimensional nutrition, and allows for physiological and comparative studies to be performed in a consistent and reproducible way from which insights onto the evolution of animal nutrition can be gained across the tree of life.

## Material and Methods

### Nutrigonometry

Studies using GF define the food components (typically macro-nutrients) that will be investigated, which together compose the ‘nutritional space’. For example, in studies where protein and carbohydrate effects are investigated, there is a 2D nutritional space (one dimension for each nutrient) onto which the performance landscape of the trait is mapped. This rationale can be extended to *n* number of nutrients (Simpson and Raubenheimer 1993*b*), albeit to date, studies with two nutrients are the most common (Morimoto and Lihoreau 2020). If we consider this 2D nutritional space as a rectangular space in which an infinite number of nutritional rails (i.e., imaginary lines that pass through the origin with arbitrary positive slope) exist that divides the space in right-angle triangles, then it is possible to use simple trigonometric functions to estimate the angle *α_i_* and the hypothenuse of the triangle, for all fitness traits mapped onto the nutritional space. The angle *α_i_* is the angle of the nutritional rail, relative from the x-axis, that passes through the peak in the landscape for the trait *i*, and the hypothenuse *h_i_* of the triangle shows how far from the origin the peak in the landscape sits for the trait *i* (Fig 1a). *α_i_* and *h_i_* can be calculated using Pythagorean theorem and the relationship between the angle and the sides of right-angle triangles (i.e., sines and cosines), as shown in Fig 1a.

**Figure 1.**
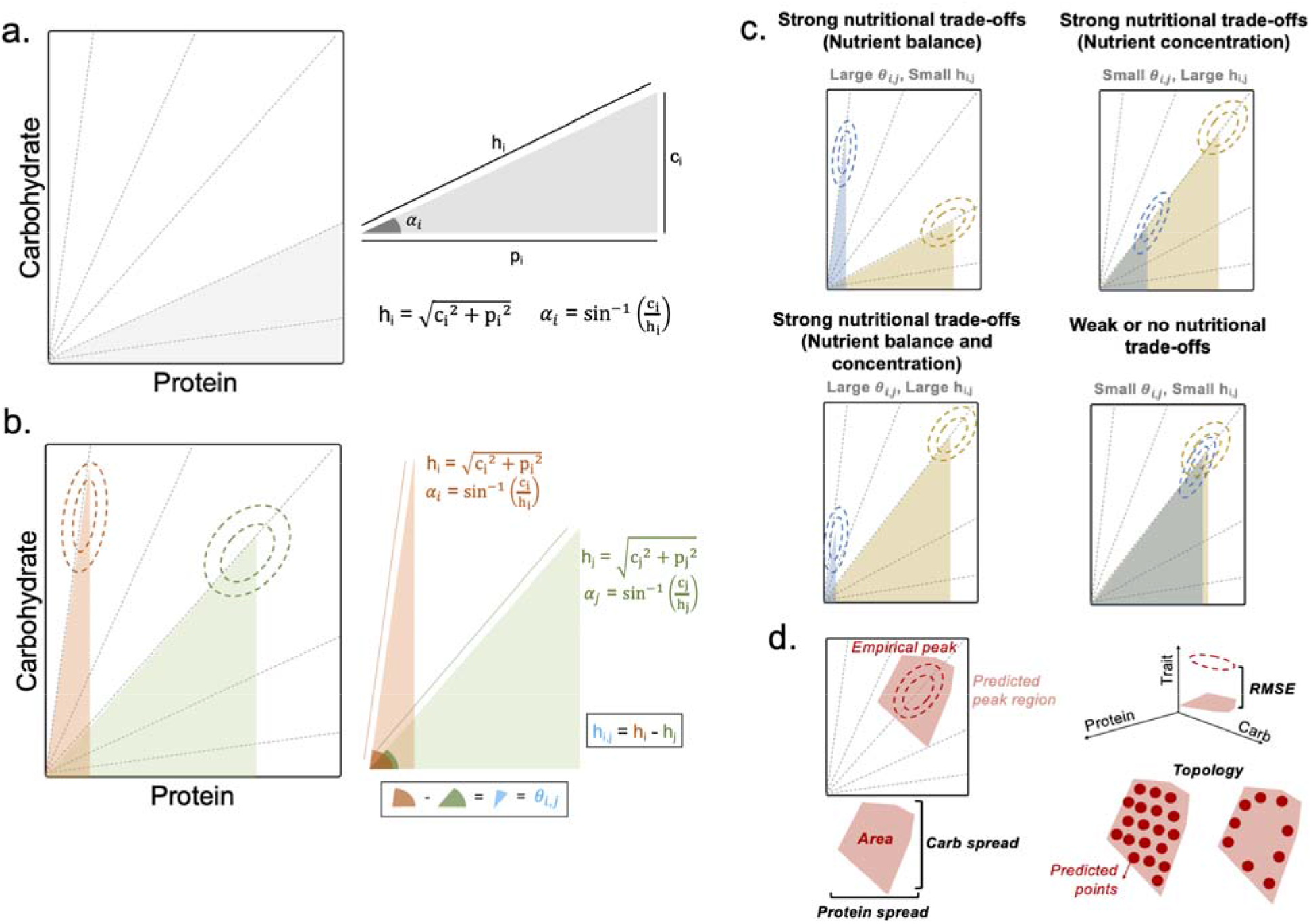
Nutrigonometry framework. (a) Considering an infinite number of nutritional rails that divide the nutritional space into right-angle triangles, the angle and the hypothenuse can be calculated from trigonometric relationships. (b) Nutrigonometry applied to compare two traits is simple as it allows for the estimates of the strength of nutritional trade-offs in terms of nutrient balance (angle) and nutrient concentration (the difference, given in absolute terms). (c) Plausible scenarios for the estimates of and . (d) Metrics used to the peak prediction in the 3D landscape. RMSE was calculated as root-mean-square difference between the predicted and observed values of the trait (z-axis) in the peak region. Nutrient spread (both carbohydrate and protein) was calculated as the standard deviation of the predicted peak region. The area of the polygon that encapsulates the predicted peak region was also estimated as a proxy of prediction performance. Lastly, the topology of the predicted datapoints for the peak region was analysed using the concept of persistence homology (see Methods) to identify homogeneity in predicted point structure.

Once *α_i_* are known, we can estimate the angle *θ* [as in (Morimoto and Lihoreau 2019)] which is the difference in the angle between nutritional rails that maximize two traits, *i* and *j*, and provides a measure of the strength of the nutritional trade-off that exists between traits *i* and *j* (Fig 1b). The larger the angle *θ_i,j_*, the strongest the nutritional trade-off in terms of nutrient balance (and potentially nutritional compromise) between traits. Likewise, we can compare the difference *h_i,j_* in the estimates of the hypothenuse *h_j_* and *h_i_* to quantify nutritional trade-offs in relation to nutrient concentration (Fig 1b). These metrics allowed us to disentangle the following scenarios in which nutritional trade-off can occur:

I. When *θ_i,J_* is large but *h_i,j_* is small (‘Strong nutritional trade-off in terms of nutrient balance’)
II. When *θ_i,J_* is small but *h_i,j_* is large (‘Strong nutritional trade-off in terms of nutrient concentration’),
III. When *θ_i,J_* and *h_i,j_* are large (‘Strong nutritional trade-off in terms of both nutrient balance and concentration’)
IV. When *θ_i,J_* and *h_i,j_* are small (‘Weak or no nutritional trade-off’) (Fig 1c).

Here when applying this model for empirical datasets (see below), inferences on the strength of nutritional trade-offs were made using confidence intervals for *h_i,j_* and *θ_i,J_*, whereby nutritional trade-offs were stronger when confidence intervals did not overlap zero and the magnitude of the difference was large. Estimates are presented in the units of the nutrient space in which the data was collected (e.g., mg), while angles are presented in degrees. Confidence intervals for both *h_i,j_* and *θ_i,J_* were calculated using the significance threshold of 0.05 and the quartiles of a *t*-distribution. All analyses and plots were done in R version 3.6.2 (R Core Team 2019).

### Predicting peak (or valley) location and size

As with previous approaches, the model presented here depends on accurate estimates of the coordinates for the peak in the multidimensional performance landscape. Without this, estimates of *h_i,j_* and *θ* are inaccurate which in turn affects the ability of the model to estimate the strength of nutritional trade-offs. To overcome this, the basic algorithms underpinning the identification of peak regions in multidimensional fitness landscapes were designed as following:

1. Empirical data was split into training (75%) and test (25%) datasets;
2. A machine learning model was fitted to the training set using 10-fold cross-validation, with the fitness trait as dependent variable and the nutrient intakes (or fixed ratios) as independent variables. The model included main and interactive effects of protein and carbohydrate, as well as quadratic effects of each nutrient (for non-linear relationships);
3. The model’s predictive performance was evaluated with root-mean-square-error (RMSE) with respect to the observed values of the test dataset;
4. A set of 500 random points corresponding to (protein, carbohydrate) coordinates were generated covering the nutritional space, and the model of step 2 was used to predict the value of the fitness value for each point;
5. A quantile threshold was used to crop the data to the specific region of interest. For instance, for peaks in the nutritional landscape, the default value used throughout this study was set to 0.95 (i.e., the highest 5% predicted fitness values are subset, from which coordinates of protein and carbohydrate are used).
6. Steps 4-5 were repeated 100 times.

We then made statistical inferences on peak area from 95% confidence intervals using the ‘ci’ function of the ‘Rmisc’ package (Hope et al. 2013) whereby we resampled with replacement the selected random points obtained from steps 5 and 6 above. To test the performance of nutrigonometry in estimating nutritional trade-offs, we used the most commonly used models to test relationships between traits in behavioral ecology (e.g., general linear model), machine learning models used in regression models in ecology and evolution [e.g., Random Forest, (Rabinovich 2021)], as well as models that have been specifically used to analyse multidimensional performance landscapes in GF studies (e.g., SVM, GAMs) (Ponton et al. 2015; Morimoto and Lihoreau 2019). In particular, we tested the performance of Bayesian linear regression (Bayes), general linear regression (LM), k-nearest neighbors (KNN), Gradient boost (GBoost), random forest (RF), support vector machine (SVM) with radial basis function as well as generalized additive models (GAMs) with both smooth term or tensor product term. With the exception of GAMs that were fitted using the ‘mgcv’ package (Wood and Wood 2015), all other models were fitted using the ‘tidymodels’ package of the tidyverse (Wickham et al. 2019). For the Bayesian regression, we used the flexible Cauchy prior from the ‘rstanarm’ package for all analysis (Goodrich et al. 2020). Fitness landscapes were estimated using the ‘Tps’ function of the ‘fields’ package (Nychka et al. 2017). All plots were done using the ‘ggplot2’ package (Wickham 2016). All models were fitted to a training set (75% of the data) and model performance (i.e., RMSE) was calculated from the performance of the models in the remaining test dataset (25%).

### Goodness-of-estimate of the models

In addition to RSME, we estimated the area (in squared units in which the data is collected) of the polygon delimited by the estimated predicted peak region (‘Area’) and the horizontal (protein) and vertical (carbohydrate) spread of the datapoints of the predicted peak region (‘Nutrient spread’) as proxies of the goodness-of-estimate of the models (Fig 1d). The smaller the RMSE, the better is the model in predicting the fitness value of the peak region (the z-axis). Furthermore, the smaller the area and nutrient spread, the more compact the prediction of the peak region in the nutritional space. Note that RMSE values do not interfere with accuracy of estimates of *h_i,j_* and *θ*, and thus the estimates of nutritional trade-offs, because the z-axis is not used in the calculation of angles and hypothenuses (Fig 1d). A model can have high RMSE and still be the best predictive model as long as the predicted peak correctly matches with the observed peak in the landscape.

### Topological structure of the estimated peak

Even in cases where area and nutrient spread of the predicted peak region are small, it is important to have evenly-spaced datapoints within the predicted peak region. This is because predictions of regions which contain holes can lead to mis-estimation of the strength of nutritional trade-offs by potentially adding noise to the set of protein and carbohydrate coordinates used to calculate *h_i,j_* and *θ*. We measured the topological structure of the predicted peak region using the concept of persistence homology (PH), which in simple terms, allows us to investigate the overall structural organisation of the data [see Text S1 and (Zomorodian and Carlsson 2005; Weinberger 2011) for details of the concept] (Fig 1d). PH was estimated using the ‘TDAstats’ package (Wadhwa et al. 2018). Together, the estimates of RMSE, area, nutrient spread, and PH provided a comprehensive suite of metrics to assess the quality of model predictions for the peak region in fitness landscapes.

### Datasets used for model application

We demonstrate the applications of the Nutrigonometry framework using two datasets, which vary in structure. The first dataset is a landmark dataset which contains *Drosophila melanogaster* individual adult nutrient intake as well as the consequences of nutrient intake to lifespan and reproduction (Lee et al. 2008). This dataset was previously used to test the Vector of Position approach and therefore has important benchmark status in the field (Morimoto and Lihoreau 2019). We also investigate the effect of data structure on Nutrigonometry estimates of peak and valley regions. In GF, data can be divided into two structures: intake data and fixed ratio data. Intake data is ideal in GF studies because it allows for exploration of *realized* nutritional effects, that is, nutritional effects exerted upon traits given by the amount of nutrient eaten (Simpson and Raubenheimer 1993*b*). However, collecting intake data can be cumbersome or challenging, and recent approaches have adapted GF experiments to draw landscapes of traits based on the fixed ratio of the nutrients in the diets (Kutz et al. 2019). To date, however, we still do not know how this adaptation influence estimates of nutritional trade-offs in multidimensional performance landscapes. Here, to test whether the structure of the data is important for model predictions, we used Lee et al., (2008) dataset with individual intakes (‘intakes’) as well as with fixed ratios (‘fixed’). Using this data and for the purpose of the demonstration of Nutrigonometry, we estimated the nutritional trade-off between lifespan and reproductive rate which are known to trade-off in this species. We also used a second dataset from (Kutz et al. 2019), which studied how temperature modulates nutritional responses of larval development and adult fitness in *D. melanogaster*, as an additional proof-of-application of our model in fixed ratio datasets (see Fig S1). Lastly, we demonstrated how the best performing models in our peak analysis can be used to predict valley regions (Fig S2 and Fig S3).

### Comparison with intake target

*Drosophila melanogaster* balance their nutrient intake to a P:C ratio of 1:4 when given the possibility to self-select multiple nutritionally complementary foods (Lee et al. 2008). We then used the peak predictions of the Nutrigonometry framework to test whether the observed P:C ratios that maximized lifespan and reproductive rates coincided with the P:C ratio of 1:4 reached by flies in choice situations. To achieve this, we calculated the 95% confidence interval as described for the peak area but in this case, for the P:C ratio of each trait. Whenever the confidence interval overlapped 1:4, we inferred that the estimate of peak ratio did not statistically differ from the intake target of 1:4.

## Results

### Simple (Bayesian) linear regressions outshine machine learning models when predicting peak region in multidimensional landscapes

All models generated predictions of peak region in nutritional landscapes irrespective of data structure although the accuracy and topology of the predicted regions varied (Fig 2 and Fig 3). In general, GAMs with tensor product and smooth function as well as Bayes and LM linear models generated peak predictions for both lifespan and reproductive rate that were significantly more accurate (narrower in area) than other models when the structure of the data was composed of food intakes (Fig 2, Table S1 and Table S2). When the data structure changed to fixed ratios, LM, GAM with tensor product, Bayes, and KNN predicted peaks with smaller area for lifespan and all but KNN perform within similar scales for the peak prediction of reproductive rate (Fig 2, Table S1 and Table S2). In comparison, GAM smooth did not perform well in predicting peak region that was homogenous and accurate to the performance landscape, particularly when the data structure was fixed ratio. The performance of the models was independent of patterns in the estimates of RMSE and nutrient spread which showed no clear pattern of performance with the exception of LM and Bayes that displayed consistently lower spread when the structure of the data were intakes (Fig 3 and Fig 4; Table S2). Interestingly, machine learning models consistently underperformed, predicting peak regions that were wider and less accurate and homogenous (Fig 2 and 3, Table S1 and S2). The underlying reason for this is unclear, but similar patterns were observed when predicting the peak region of (Kutz et al. 2019) dataset (see Fig S1 and Table S3). Bayes, GAMs (both smooth and tensor product) and LM also performed well when predicting valley regions (see Fig S2 and S3). These results indicate that simple (Bayesian) linear regression provide consistently the best models to estimate the region of the peak of fitness landscapes irrespective of the structure of the data, and that GAMs with tensor product (and to a smaller extent, smooth function) can be used when the data are individual intakes.

**Figure 2.**
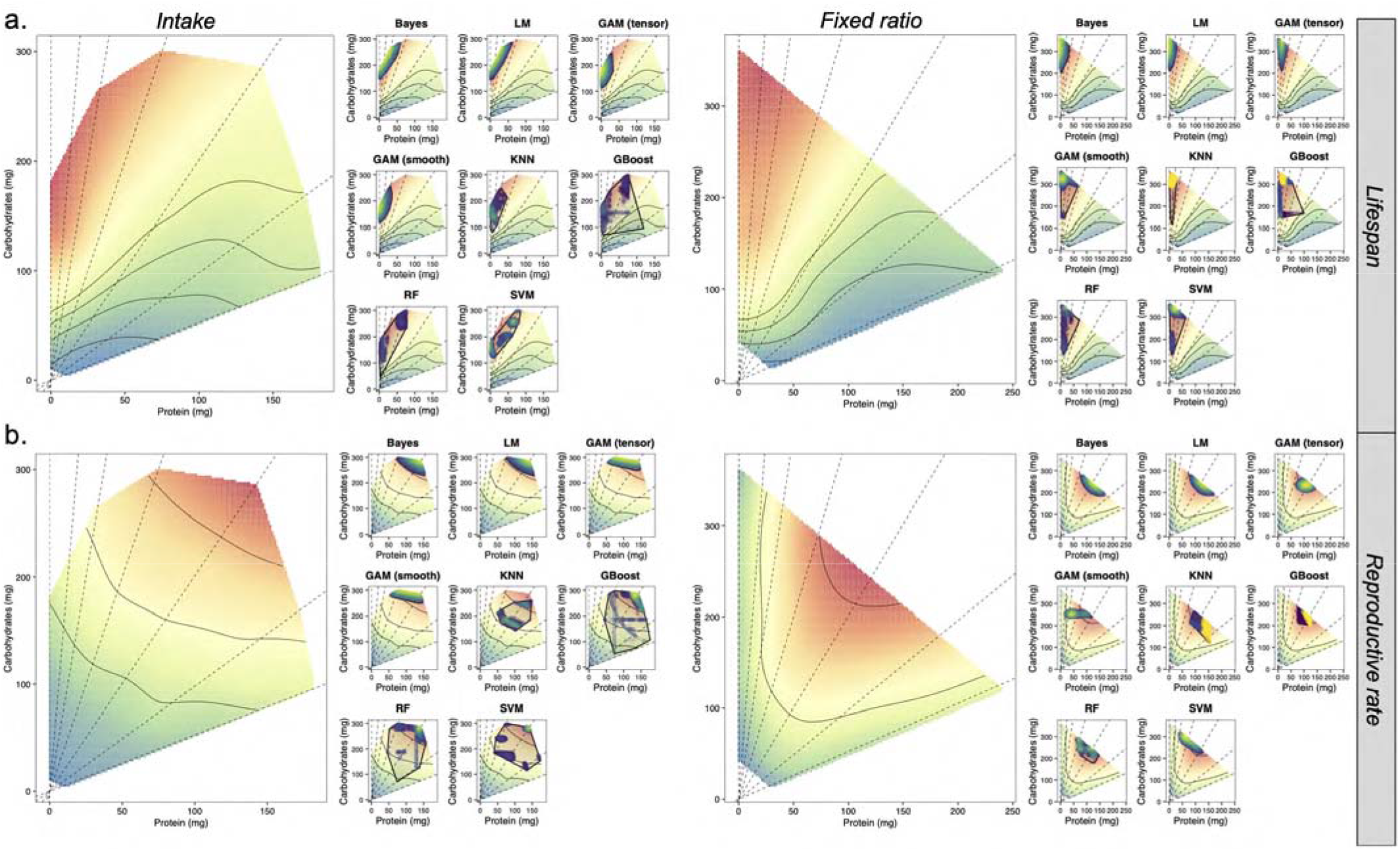
Nutrigonometry framework to predict peak region in lifespan and reproductive rate landscape with different data structure. (a) Lifespan landscapes with individual intake (top left panel) and fixed ratio (top right panel) from Lee et al. (2008) with the overlaid predicted peak regions. (b) Reproductive rate landscapes with individual intake (bottom left panel) and fixed ratio (bottom right panel) from Lee et al. (2008) with the overlaid predicted peak regions. For the landscapes, red represents peaks while light green represents valleys. For the predicted region, dark blue represents points with lower predicted z-values whereas bright yellow represents points with higher predicted z-values. The shaded polygon was added to facilitate visualisation of the predicted peak region and the homogeneity of points within the predicted peak.

**Figure 3.**
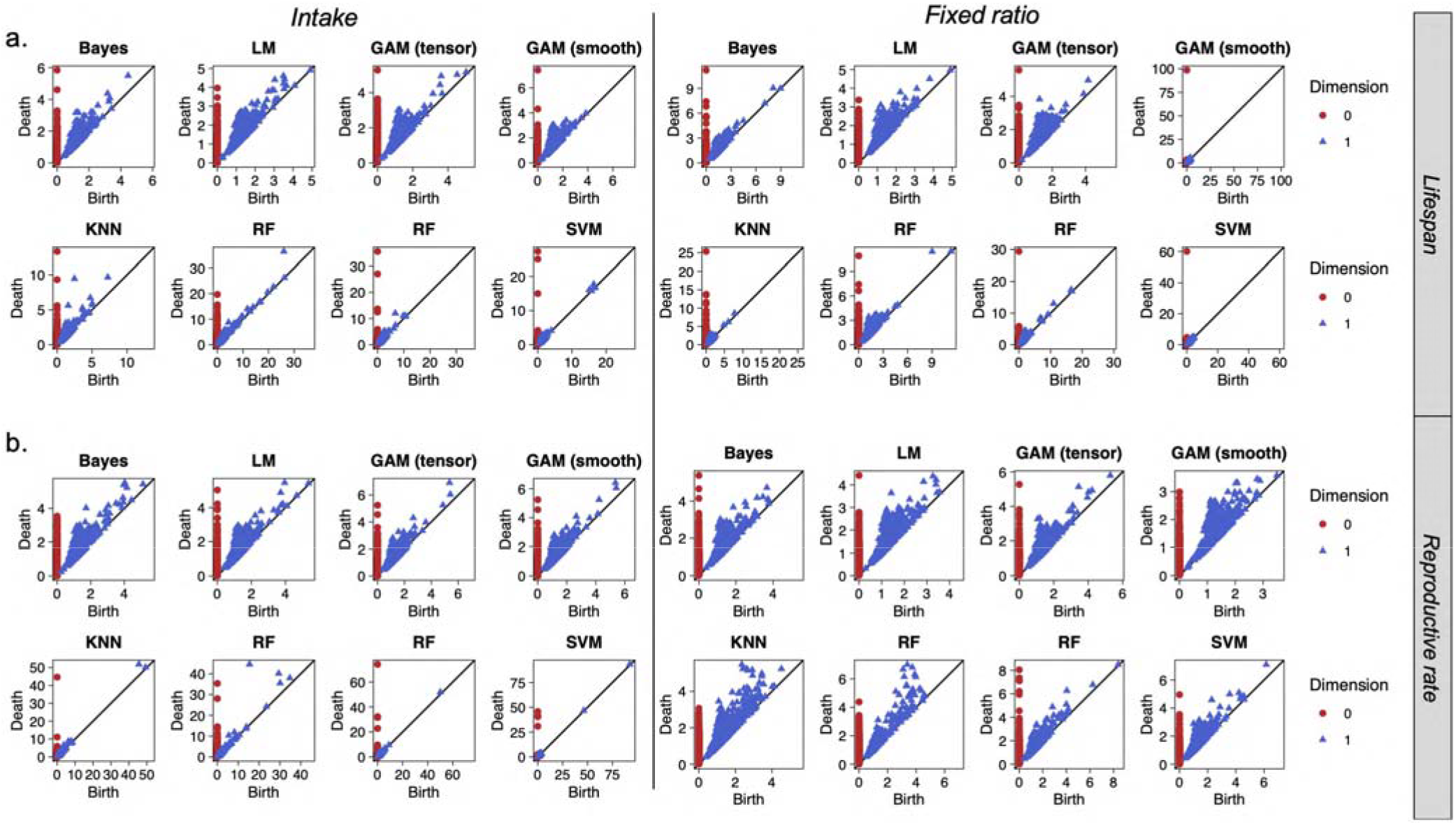
Persistence Homology (PH) to investigate topological structure of the predicted peak region using Nutrigonometry. (a) PH plots for the topological analysis of the predicted peak region in lifespan of data containing the structure of individual intake (top left) and fixed intake data (top right). (b) PH plots for the topological analysis of the predicted peak region in reproductive rate with data of structure containing individual intake (bottom left) and fixed intake data (bottom right). Homogenous predicted peaks have red (dimension 0) and blue (dimension 1) points that are closer, as opposed to more heterogeneous predicted peaks upon which (some) points can be farther from each other. Note the different scales upon which the data is plotted (needed to aid visualisation of point clouds).

**Figure 4.**
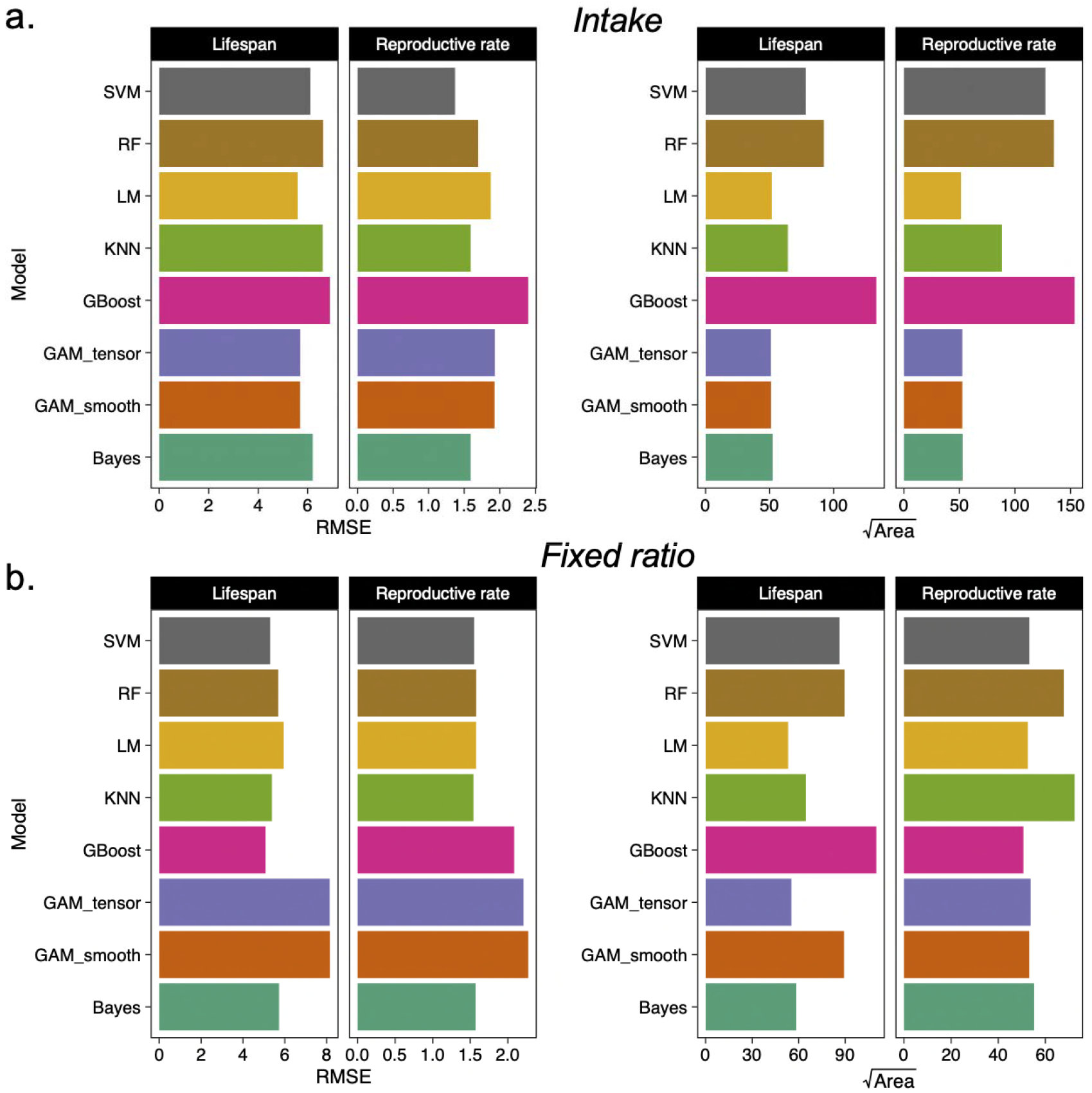
RMSE and peak area estimates in the Nutrigonometry peak region predictions. (a) RMSE and predicted peak area (i.e., area of the shaded polygon from the predicted region for lifespan and reproductive rate data), with structure containing individual intakes. (b) RMSE and predicted peak area (i.e., area of the shaded polygon from the predicted region for lifespan and reproductive rate data), with structure containing fixed ratios. Note that models with high RMSE can still be the best predictors of peak region.

### Better models lead to accurate estimates of known nutritional trade-offs in multidimensional landscapes

GAMs (both smooth and tensor product), Bayes, LM, and KNN models were the only models that correctly identified the nutritional trade-off measured by 0 between lifespan and reproductive rate for data with individual intakes (Table 1). Given the variability in the area, spread, and topology of the predicted region, estimates of *h_i,j_* and *θ* were more accurate (narrower confidence intervals) for GAMs (smooth and tenor product), Bayes and LM compared with KNN. GAMs, Bayes and LM were the only ones that identified a trade-off on the hypothenuse estimate *h_i,j_* for data of individual intakes, while KNN was the only model that identified this trade-off in data with fixed ratio. GAM smooth was the only model that failed to identify this trade-off when the data was composed of fixed ratios (Table 1). Thus, overall, simpler models are more suitable to generate peak predictions that accurately describe nutritional trade-offs in multidimensional performance landscapes for data of different structures.

**Table 1.** Nutrigonometry quantification of nutritional trade-offs between lifespan and reproduction. Estimates of *θ_i,J_* (in degrees) and *h_i,j_* (in mg) for the nutritional trade-off between lifespan and reproductive rate. Analysis from the data presented in Lee et al. (2008). Confidence intervals overlapping zero implies no difference in the peaks. Magnitude of the estimates indicate the strength of nutritional trade-offs (i.e., larger magnitudes indicate stronger nutritional trade-offs). Note that *θ_i,J_* is bound between 0 and 90 degrees (i.e., 0 and 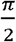).

|  | Parameter | Model | Estimate | Std | LwrCI | UprCI |
| --- | --- | --- | --- | --- | --- | --- |
| Trade-off<br>(intakes) | $\theta_{i,j}$ | SVM | 14.456 | 10.728 | -6.574 | 35.485 |
|  |  | RF | 14.508 | 8.109 | -1.388 | 30.404 |
|  |  | GAM_tensor | 16.128 | 4.984 | 6.358 | 25.897 |
|  |  | GAM_smooth | 16.166 | 4.962 | 6.438 | 25.893 |
|  |  | GBoost | 17.063 | 9.575 | -1.706 | 35.831 |
|  |  | LM | 17.940 | 4.826 | 8.479 | 27.400 |
|  |  | Bayes | 18.205 | 4.709 | 8.974 | 27.436 |
|  |  | KNN | 21.203 | 6.181 | 9.088 | 33.318 |
| | $h_{i,j}$ | SVM | 16.792 | 65.723 | -112.038 | 145.622 |
|  |  | KNN | 50.015 | 48.137 | -44.343 | 144.373 |
|  |  | GBoost | 52.851 | 75.218 | -94.591 | 200.293 |
|  |  | RF | 58.561 | 66.066 | -70.943 | 188.064 |
|  |  | LM | 75.870 | 35.142 | 6.984 | 144.757 |
|  |  | Bayes | 76.729 | 34.444 | 9.211 | 144.247 |
|  |  | GAM_smooth | 120.245 | 29.406 | 62.604 | 177.886 |
|  |  | GAM_tensor | 124.533 | 27.930 | 69.784 | 179.282 |
| Trade-off<br>(fixed) | $\theta_{i,j}$ | GAM_smooth | 9.645 | 5.897 | -1.916 | 21.205 |
|  |  | SVM | 11.840 | 5.649 | 0.767 | 22.913 |
|  |  | RF | 17.368 | 5.848 | 5.906 | 28.831 |
|  |  | GBoost | 20.177 | 5.057 | 10.264 | 30.090 |
|  |  | GAM_tensor | 21.177 | 3.872 | 13.588 | 28.766 |
|  |  | Bayes | 26.454 | 5.876 | 14.935 | 37.973 |
|  |  | LM | 26.499 | 5.903 | 14.928 | 38.070 |
|  |  | KNN | 31.428 | 7.186 | 17.342 | 45.513 |
| | $h_{i,j}$ | SVM | 2.381 | 68.888 | -132.653 | 137.416 |
|  |  | RF | 4.819 | 64.841 | -122.283 | 131.921 |
|  |  | GBoost | 9.377 | 65.605 | -119.222 | 137.975 |
|  |  | Bayes | 41.461 | 34.912 | -26.974 | 109.896 |
|  |  | LM | 42.305 | 34.429 | -25.182 | 109.791 |
|  |  | GAM_smooth | 46.635 | 40.358 | -32.475 | 125.745 |
|  |  | GAM_tensor | 49.009 | 32.855 | -15.394 | 113.412 |
|  |  | KNN | 82.516 | 30.388 | 22.949 | 142.083 |

### Comparing trait optimum with intake target

We then used the estimated peak regions for lifespan and reproductive rate (both individual intake and fixed ratio data structures) to estimate the optimum P:C ratio that maximises each trait as well as whether or not these optima coincided with the P:C ratio obtained when individuals are allowed to balance their diet (i.e., 1:4). All models predicted a significantly lower P:C ratio for the optimum that maximizes reproductive rate relative to lifespan as expected from the original visual comparison of landscapes (around 1:2 for reproductive rate and >1:9 for lifespan) (Table 2). However, none of the estimates overlapped 1:4, suggesting that *D. melanogaster* females likely have to compromise the nutrient intake to maximise either lifespan or reproductive rate, but not both simultaneously.

**Table 2.** Nutrigonometry estimates of nutritional compromises. Estimates of optimal intake that maximises lifespan and reproductive rate based on the predicted peak region. Comparison made with the visual peak ratio from Lee et al. (2008). Note that all models show that the estimated peak ratio between traits do not overlap and thus, corroborate the inference of a nutritional trade-off between traits, leading to a nutritional compromise. Note also that all but one model (i.e., GAM smooth for fixed ratio reproductive rate data) predicted peak region ~1:4, which is the ratio that individuals balance when given the ability to balance their diet (‘choice’). All other models suggest that a P:C ratio of 1:4 is lower than the ratio needed to maximise lifespan but higher than that for reproductive rate, further supporting the concept of a nutritional compromise.

| Data | Trait | Model | Mean | Upr CI | Lwr CI | Target (Visual) |
| --- | --- | --- | --- | --- | --- | --- |
| Peak<br>Ratio (intakes) | Lifespan | GBoost | 5.235 | 5.205 | 5.265 | 16 |
|  |  | RF | 5.533 | 5.508 | 5.557 |  |
|  |  | SVM | 5.864 | 5.836 | 5.892 |  |
|  |  | LM | 9.084 | 9.048 | 9.120 |  |
|  |  | Bayes | 9.154 | 9.118 | 9.190 |  |
|  |  | KNN | 12.075 | 12.015 | 12.135 |  |
|  |  | GAM_smooth | 13.055 | 12.997 | 13.114 |  |
|  |  | GAM_tensor | 13.108 | 13.049 | 13.168 |  |
|  | Reproductive<br>rate | GBoost | 1.858 | 1.853 | 1.864 | 2 |
|  |  | KNN | 2.041 | 2.037 | 2.045 |  |
|  |  | RF | 2.138 | 2.133 | 2.144 |  |
|  |  | SVM | 2.147 | 2.138 | 2.156 |  |
|  |  | Bayes | 2.194 | 2.191 | 2.197 |  |
|  |  | LM | 2.215 | 2.212 | 2.219 |  |
|  |  | GAM_smooth | 2.644 | 2.639 | 2.650 |  |
|  |  | GAM_tensor | 2.661 | 2.656 | 2.667 |  |
| Peak<br>Ratio (fixed) | Lifespan | GBoost | 13.078 | 12.987 | 13.170 | 16 |
|  |  | SVM | 14.946 | 14.873 | 15.019 |  |
|  |  | RF | 15.107 | 15.045 | 15.169 |  |
|  |  | GAM_smooth | 16.977 | 16.859 | 17.097 |  |
|  |  | GAM_tensor | 20.623 | 20.536 | 20.710 |  |
|  |  | KNN | 26.050 | 25.929 | 26.173 |  |
|  |  | Bayes | 32.426 | 32.237 | 32.617 |  |
|  |  | LM | 32.933 | 32.744 | 33.125 |  |
|  | Reproductive<br>rate | KNN | 1.467 | 1.463 | 1.470 | 2 |
|  |  | LM | 1.836 | 1.832 | 1.839 |  |
|  |  | Bayes | 1.841 | 1.837 | 1.845 |  |
|  |  | GBoost | 2.171 | 2.168 | 2.174 |  |
|  |  | GAM_tensor | 2.244 | 2.241 | 2.248 |  |
|  |  | RF | 2.545 | 2.539 | 2.552 |  |
|  |  | SVM | 3.527 | 3.516 | 3.538 |  |
|  |  | GAM_smooth | 4.280 | 4.265 | 4.296 |  |

## Discussion

We proposed a new simple analytical framework to analyse nutritional trade-offs in multidimensional fitness landscapes. Nutrigonometry uses trigonometric relationships from right-angle triangles to identify and compare peaks (or valleys) in 3D fitness landscapes between traits. Using landmark GF datasets with different structures, we demonstrated the accuracy and performance of standard (machine learning) models in finding the peak regions in these multidimensional landscapes and subsequently quantifying the strength of nutritional trade-offs between traits. As with the Vector of Positions approach (Morimoto and Lihoreau 2019), the Nutrigonometry strictly considers coordinates in the real positive region of the nutrient space, whereby the true separation between key regions (peaks and valleys) are quantified within the correct domain in which the fitness landscapes exist. However, contrary to previous methods (Rapkin et al. 2018; Morimoto and Lihoreau 2019), the Nutrigonometry does not rely on vector calculations but instead harnesses the trigonometric relationships of right-angle triangles to estimate nutritional trade-offs. This is a major advance of the model as it considerably simplifies the framework both in conceptual and computational terms. Nutrigonometry thus significantly advances our ability to generate reliable and reproducible estimates of nutritional trade-offs within and between species, facilitating quantitative (comparative studies) of animal nutrition.

Multidimensional studies of nutrition through the GF have been increasingly used to gain insight into animal and human nutrition (Lee et al. 2008; Behmer 2009; Felton et al. 2009; Simpson and Raubenheimer 2012; Hewson-Hughes et al. 2013; Gosby et al. 2014; Solon-Biet et al. 2014). Likewise, the complexity of the applications has also increased, ranging from studies with few nutrients (e.g., protein and carbohydrates, salts) through to high-dimensional studies investigating individual fatty acids and amino acids (Simpson et al. 2006; Grandison et al. 2009; Arien et al. 2015; Arganda et al. 2017; Piper et al. 2017). This means that analytical frameworks that are simple and robust must be developed to support the development of the field. Nutrigonometry provides such foundation, by demonstrating the best approach to investigate nutritional trade-offs in 3D fitness landscapes. Because Nutrigonometry uses trigonometric relationships of right-angle triangle, it is applicable to *n* dimensions. However, given the often counter-intuitive geometrical effects of high dimensionality [e.g., (Milman 1998; Watanabe 2021)], such expansion to higher dimensions requires further investigation as the topic of future developments. Nevertheless, given the broad use of 3D fitness landscapes in GF studies (Morimoto and Lihoreau 2020), Nutrigonometry readily enables important quantifications of nutritional trade-offs that were otherwise absent or cumbersome to produce. For instance, using a range of models, Nutrigonometry uses right-angle triangles to compare the ratio of nutrients that maximise lifespan and reproductive rate along with the strength of nutritional trade-offs between these traits in a landmark paper in the field (Lee et al. 2008). Moreover, Nutrigonometry is capable of comparing the nutrient ratio which maximises lifespan and reproductive rate with the nutrient ratio that is balanced by individuals when given a choice, providing important insights into the dietary choices underpinning nutritional compromises. Such quantification can bring new fundamental insights into our understanding of nutritional trade-offs such as strength and the direction of the trade-offs (e.g. nutrient balance vs concentration, see Fig. 1), as well as how much animals actually resolve these trade-offs when they have the opportunity to do so and whether, for instance, they favour one trait of another (distance between optimal trade-off and observed nutrient intake target, see Table 2).

An important trend in the field of multidimensional nutrition is the study of nutritional effects across physiological pathways and across levels of biological organization (Lihoreau et al. 2014; Simpson et al. 2015). These studies generate multiple 3D landscapes that are often compared visually, without rigorous analytical methods to measure nutritional trade-offs. For example, eleven 3D-landscapes of the expression of genes involved in the Insulin/IGF pathway were visually compared to provide insights into how a key endocrine pathway is regulated based on nutrient intake, and how gene expression can underlie expression of life-histories (Post and Tatar 2016; McDonald et al. 2021). Likewise, twelve 3D-landscapes with gut microbial diversity or abundance were visually compared to better understand how nutrient composition can modulate host-microbe interactions (Ng et al. 2019). Similar visual comparisons have been made to understand the effects of nutrition on host-endosymbiont relationship (Ponton et al. 2015). The analytical framework proposed here will allow researchers to move beyond visual comparisons to quantitatively assess how landscapes differ using a rigorous and reproducible framework. As a result, Nutrigonometry yields considerable advances to the status quo in the field, enabling a deeper understanding of the role of nutrition in host-microbial interactions as well as animal physiology, behaviour and ecology.

## Supporting information

ESM

## Acknowledgements

ML receives support from the CNRS, the French Research Agency (ANR 3DNaviBee: ANR-19-CE37-0024), the European Regional Development Fund (FEDER ECONECT: MP0021763), and the European Research Council (ERC-CoG BEE-MOVE: GA101002644).

## Data accessibility

Kutz et al. data is available here: doi.org/10.26180/5cfe1ddaaafac. Lee et al. dataset is available in Dryad: doi:10.5061/dryad.tp7519s. R script with functions for the implementation of the Nutrigonometry framework is available in the Supplementary Materials.

## Supplementary Material

**Text S1. What is Persistence Homology (PH)? A brief introduction.**

**Table S1. Area of the predicted peak region for all models**. All values are given in unit squared of nutrient intake or diet composition (for fixed ratios).

**Table S2. Nutrient spread of the predicted peak region for all models**. All values are given in units of nutrient intake or diet composition (for fixed ratios).

**Table S3. Nutrigonometry quantification of nutritional trade-offs in developmental time between two developmental temperatures**. Estimates of *θ_i,J_* (in degrees) and *h_i,j_* (in g/L). Analysis from the data presented in Kutz et al. (2018). Confidence intervals overlapping zero implies no difference in the peaks. Magnitude of the estimates indicate the strength of nutritional trade-offs (i.e., larger magnitudes indicate stronger nutritional trade-offs). Note that *θ_i,J_* is bound between 0 and 90 degrees (i.e., 0 and 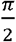).

**Figure S1**. (a) 3D landscape for developmental time at 25°C (top left) and 28°C (bottom left) (from Kutz et al., 2019) with the overlaid predicted peak regions. For the landscape, red represents peaks while light green represents valleys. For the predicted region, dark blue points represent points with lower predicted z-values whereas bright yellow represents points with higher predicted z-values. The shaded polygon was added to facilitate visualisation of the predicted peak region and the homogeneity of points within the predicted peak. (b) RMSE and predicted peak area (i.e., area of the shaded polygon in panel *a*) for the models of developmental time at 25°C (top right) and 28°C (bottom right) values of each model. Note that models with high RMSE can still be the best predictors of peak region. (c) Persistence homology (PH) plots for the topological analysis of the predicted peak region of the 3D landscape for developmental time at 25°C (top panel) and 28°C (bottom panel) (from Kutz et al., 2019). x and y-axes represent birth and death, respectively, of topological structures. The diagonal line represents the line in which the birth and death co-occur. Homogenous predicted peaks have red (dimension 0) and blue (dimension 1) points that are closer, as opposed to more heterogeneous predicted peaks upon which points are farther from each other.

**Figure S2. Prediction of the valley regions for lifespan using individual intake data from Lee et al. (2008)**. Note that we used the best performing models for the peak region (see Main text).

**Figure S3. Prediction of the valley regions for reproductive rate using individual intake data from Lee et al. (2008)**. Note that we used the best performing models for the peak region (see Main text).

**R Script**. R script with functions for the implementation of the Nutrigonometry framework (separate file).

