## Supplementary material for "Nutrigonometry I: using right-angle triangles to quantify nutritional trade-offs in multidimensional performance landscapes": ESM

Authors’ affiliations:

^1^ School of Biological Sciences, University of Aberdeen, Zoology Building, Tillydrone Ave, Aberdeen AB24 2TZ

^2^ Institute of Mathematics, University of Aberdeen, King's College, Aberdeen AB24 3FX

^3^ School of Biological Sciences, Monash, University, Melbourne, Victoria, Australia

^4^ Research Center on Animal Cognition (CRCA), Center for Integrative Biology (CBI); CNRS, University Paul Sabatier – Toulouse III, France.

* Correspondence:

Dr Juliano Morimoto

**Running-title:** Trigonometry to measure fitness trade-offs

**Keywords**: Nutritional Geometry, nutritional trade-off, performance landscapes

**Text S1 – What is Persistence Homology (PH)? A brief introduction**

***Topological data analysis and persistence homology***

Topological data analysis (TDA) is a relative new subject that lies at the intersection of algebraic topology, data science, statistics and computer science (Chazal & Michel, 2021). Algebraic topology is the part of mathematics that studies (topological) spaces by associating algebraic objects to it that do not change under ‘well-behaved’ deformations called *homotopies*, e.g., affine transformations – scaling, rotations, translations – and more generally ‘squeezing and stretching the space without tearing it apart’. Such algebraic invariants, that is, those that do not vary upon these deformations, are commonly called *topological invariants*. In a way, one could encapsulate the goal of the field of TDA as: *data has shape* and TDA aims to measure it.

One of the branches of TDA is called *persistence homology* (PH) which has provided new insights and methods to understand higher dimensional data and extra qualitative features of data using techniques of algebraic topology. In the last years, PH has been extensively studied and, by now, is supported by a solid theoretical framework to study complex data structures. Likewise, PH application is supported by a range of well-established softwares designed for PH analysis pipeline, making PH more accessible to different scientific communities. The aim of PH is to extract qualitative information of the data via the computation of a topological invariant called *homology.* For the purpose of the Nutrigonometry, it is enough to think of homology as an algebraic tool that records the number of holes on each dimension of a given space. However, what is a n-dimensional hole? A 0-dimensional hole is the number of connected components (e.g., points). A 1-dimensional hole is the number of cycles/loops that are formed when connected components are linked. A 2-dimensional hole is the number of holes enclosed by a surface (like the sphere, i.e., the boundary of a ball, for instance) and so on (see below for an example). Note that holes in a space are invariants under homotopies, in other words, a hole cannot be created or erased in homotopical deformation. This is important because it shows that the topological structure of the data and the information obtained from these n-dimensional holes are invariant. As a result, homology is then an algebraic structure associated to the input data robust under small perturbations (deformations).

The overall structure of a PH pipeline is as following:

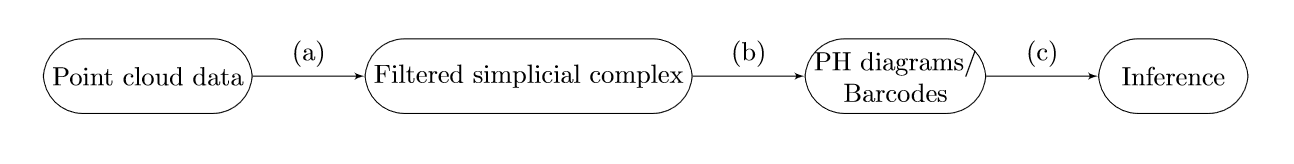

We move from the original cloud of datapoints, upon which we implement a filtering function that allows us to extract the n-dimensional holes described above (a). We then display the information of the n-dimensional holes with standard plots (b) from which inferences on the topological structure of the data can be made (c). For the purpose of this paper, we will focus on providing a user-guide explanation on how to read and interpret persistence diagrams. For a detailed survey and introduction on PH, refer to (Otter, Porter, Tillmann, Grindrod, & Harrington, 2017) and (Ghrist, 2008). For an introduction on algebraic topology and precise mathematical definitions of the concepts defined here, consult (Ghrist, 2014; Hatcher, 2002).

***From point cloud data to a realized space: filtered simplicial complex***

Topological spaces, as generalizations of geometric objects, are all around and with a rich variety of examples – from Euclidean space to fractals. It can be hard to compute the homology of a general space. However, for most of the purpose of PH and most of applications the realized space is a topological space called *simplicial complex.* A simplicial complex is built out of pieces called simplices which occur in any dimension *n*, where *n* is a non-negative integer. The 0-simplices are points, the 1-simplices are edges, the 2-simplices are triangles, the 3-simplices are tetrahedrons and so on. More precisely, a n-simplex represent a convex hull of n+1 points in the n dimensional Euclidean space that are affinely independent, that is, are not all on the same n-1 dimensional hyperplane*.*

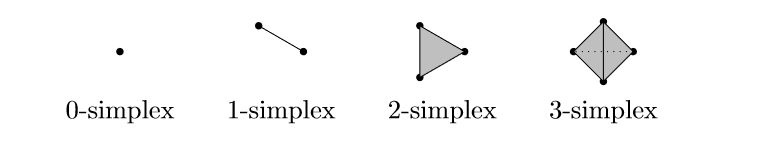

Note that a simplex is determined by its vertex set, hence standard notation of a n-simplex is σ = [v_1_, … , v_n_]. Each simplex has what is called *boundary faces,* simplices of dimension one below their own. For instance, a 1-simplex has two 0-simplices as boundary faces, a 2-simplex has three 1-simplices as boundary faces and, more generally, a n-simplex has n+1 (n-1)-simplices as boundary faces. Finally, a simplicial complex is built out of simplices by gluing them together with only one rule to be satisfied: two simplices of any dimension can be glued along a common boundary faces of the same dimension. This surprisingly naive definition has led to important developments in mathematics.

The first step is to construct a simplicial complex out of the point cloud data. One of the most natural ways to do it is via the *Vietoris-Rips complex or filtration*. Recall that our data is embedded in the Euclidean space and it makes sense to talk about (Euclidean) distance. Let ε be greater or equal than 0. The Vietoris-Rips complex for ε is the simplicial complex whose k-simplices are the k+1 data point that are pairwise ε distant. A way to visualize it is the following: at each data point we draw a ball of diameter ε, if k+1 balls intersect there is a k-simplex. For instance, for very small ε the associated Vietoris-Rips complex is a discrete set of point (the data point themselves) and for very large ε a n-simplex, where n is the number of data points. In other words, there is a filtration by varying the value of the scale parameter ε, we start with ε equals to zero until there is no more change in the simplicial complex, that is, the Vietoris-Rips complex becomes a simplex of a given dimension. More precisely, denote VR = (VR_i_)_n_ a sequence of Vietoris-Rips complexes associated to data set for an increasing sequence of scale parameter ε_i_, and we have a sequence of inclusion of topological spaces

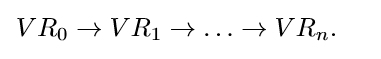

***Persistence diagrams: a topological signature of the data***

The next step is to compute the homology of the space VR = (VR_i_)_n_ at each step of the filtration, that is, the number of holes of the space at each step of the filtration according to scale parameter ε. The name persistence homology comes from the fact that we observe which parts of the homology (for each possible dimension) *persists* as the scale parameter ε_i_ increases. There are two equivalent ways of visualising the homological calculation: via persistence *diagrams* or via *barcodes*. One can think of a persistence diagram/barcode as a representation of a topological signature of the data. For sake of completeness, we present both.

A persistence diagram is a two-dimension plot where each point there represents an apparition of the topological feature in question (e.g., a hole) in the data set. The value horizontal coordinate (x-axis) indicates the **birth** time of a hole and the vertical coordinate (y-axis) is the **death** time of it according to ε_i_. The death of each hole always occurs after its birth, and thus, all the points in the persistence diagram must lie above the diagonal line. Note that points close to the diagonal line are features (hole) with a really short ‘lifespan’.

An alternative way to interpret the same information is through the barcode plot, where each bar in the barcode represents an apparition of a hole in the dataset, with the beginning of the bar showing its **birth** the end of the bar its **death** as the scale parameter varies in the filtration VR. Short bars in the barcode’s representation are equivalent to points that lie in the diagonal line in the persistence diagram, whereas long bars are point far from the diagonal line.

As an example, consider the point cloud data **A** below

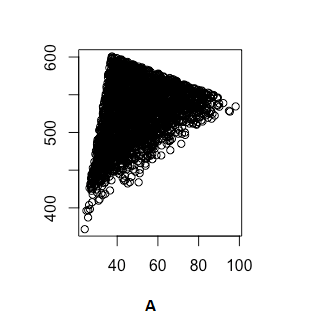

The persistence diagram (on the left) and the barcode representation (on the right) of **A** are given below

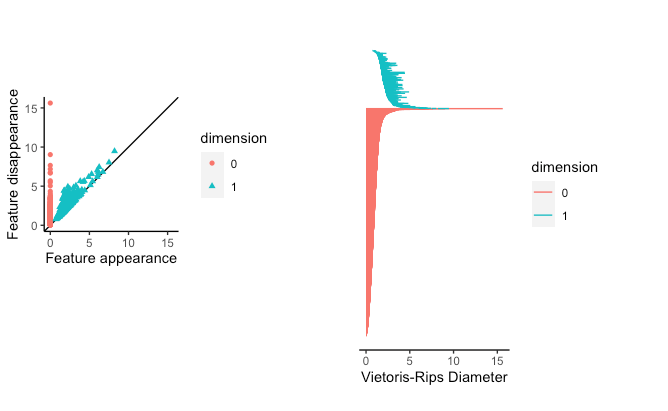

Note that all the red points (i.e., connected components or 0-dimension simplex) are aggregated and have a really short lifespan as the scale parameter grows. Moreover, the blue points (i.e., cycles or 1-dimension hole) also have a relative short life space. *This usually indicates that all the data points lie in a unique cluster with points really close to each other and relatively homogenous, since there is no space to form a cycle within a persistent enough interval with respect to the scale parameter.* This is the property used to measure homogeneity in peak (or valley) region prediction in using the Nutrigonometry (see Main Text).

***Interpretation***

As a direct and easy consequence, one is able to spot when two data sets are different by looking at their persistence diagrams. Moreover, a persistence diagram will give *a global analysis of the data*. Higher points in persistence diagrams correspond to more persistent features of the data and potentially more informative and points close to the diagonal (point with short life-span) are regarded as noise or small perturbation. However, recent studies have shown that such short-life points may contain information about *the local geometry properties* of data (Adams & Moy, 2021) and, therefore, their significance will depend on the problem/data in consideration. A current research topic to obtain a detailed statistical inference out of persistence diagrams/barcodes, as outlined in (Otter et al., 2017) and there are still challenges and open questions to be answered. Finally, PH has been used to machine learning applications as a way of providing feature vectors and enhancing machine learning algorithms.

***References cited***

Adams, H., & Moy, M. (2021). Topology Applied to Machine Learning: From Global to Local. *Frontiers in Artificial Intelligence*, *4*, 54.

Chazal, F., & Michel, B. (2021). An introduction to topological data analysis: fundamental and practical aspects for data scientists. *Frontiers in Artificial Intelligence*, *4*.

Ghrist, R. (2008). Barcodes: the persistent topology of data. *Bulletin of the American Mathematical Society*, *45*(1), 61–75.

Ghrist, R. (2014). *Elementary applied topology* (Vol. 1). Createspace Seattle, WA.

Hatcher, A. (2002). Algebraic topology, Cambridge Univ. *Press, Cambridge*.

Otter, N., Porter, M. A., Tillmann, U., Grindrod, P., & Harrington, H. A. (2017). A roadmap for the computation of persistent homology. *EPJ Data Science*, *6*, 1–38.

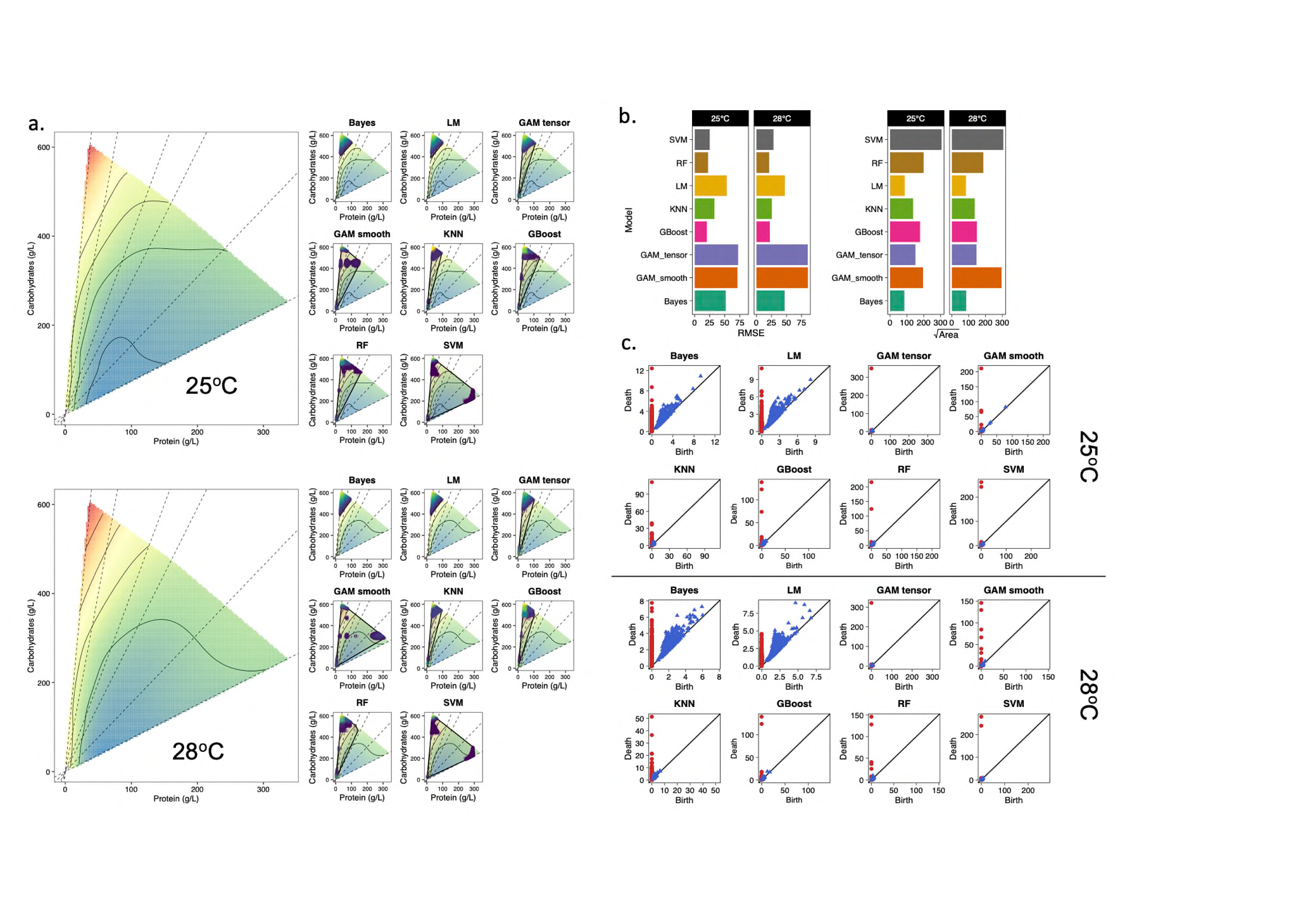
**Supplementary Figures**

**Figure S1.** (a) 3D landscape for developmental time at 25^o^C (top left) and 28^o^C (bottom left) (from Kutz et al., 2019) with the overlaid predicted peak regions. For the landscape, red represents peaks while light green represents valleys. For the predicted region, dark blue points represent points with lower predicted z-values whereas bright yellow represents points with higher predicted z-values. The shaded polygon was added to facilitate visualisation of the predicted peak region and the homogeneity of points within the predicted peak. (b) RMSE and predicted peak area (i.e., area of the shaded polygon in panel *a*) for the models of developmental time at 25^o^C (top right) and 28^o^C (bottom right) values of each model. Note that models with high RMSE can still be the best predictors of peak region. (c) Persistence homology (PH) plots for the topological analysis of the predicted peak region of the 3D landscape for developmental time at 25^o^C (top panel) and 28^o^C (bottom panel) (from Kutz et al., 2019). x and y- axes represent birth and death, respectively, of topological structures. The diagonal line represents the line in which the birth and death co-occur. Homogenous predicted peaks have red (dimension 0) and blue (dimension 1) points that are closer, as opposed to more heterogeneous predicted peaks upon which points are farther from each other.

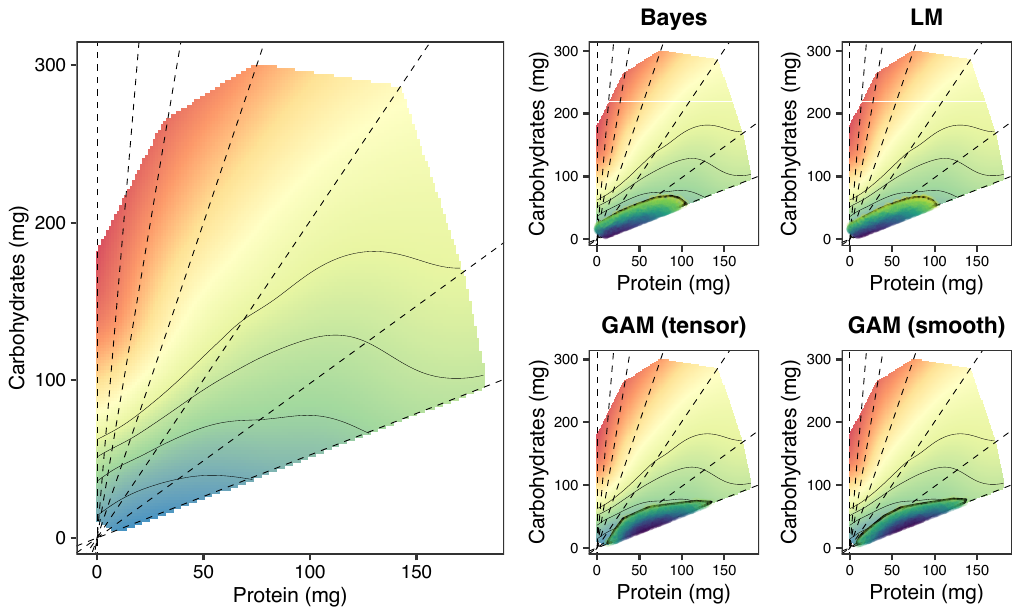

**Figure S2. Prediction of the valley regions for lifespan using individual intake data from Lee et al., 2008.** Note that we used the best performing models for the peak region (see Main text).

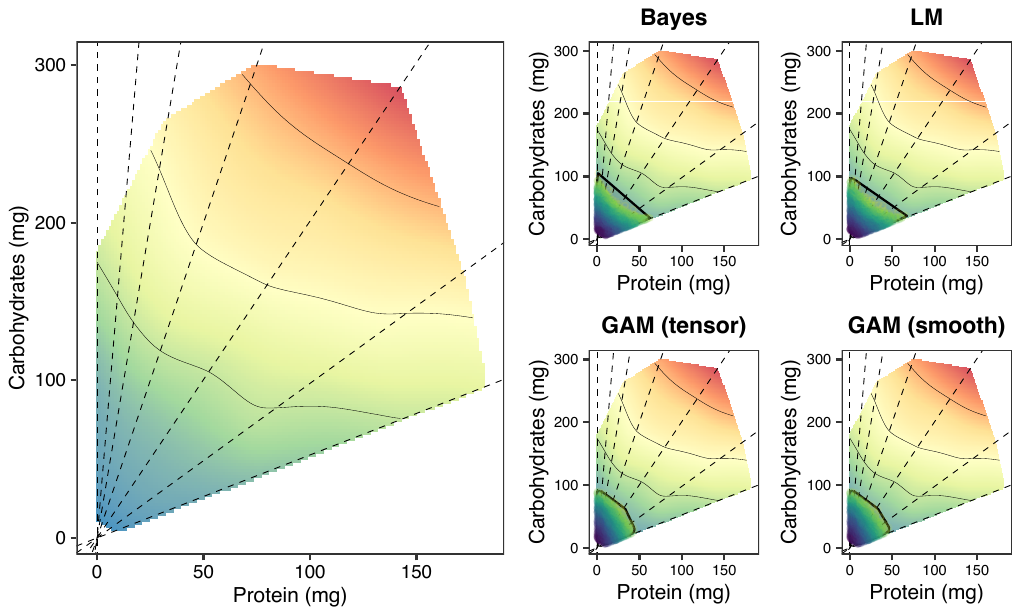

**Fig S3. Prediction of the valley regions for reproductive rate using individual intake data from Lee et al. (2008).** Note that we used the best performing models for the peak region (see Main text).

**Supplementary Tables**

**Table S1. Area of the predicted peak region for all models.** All values are given in unit squared of nutrient intake or diet composition (for fixed ratios).

| **Data** | **Trait** | **Model** | **Mean** | **Upr CI** | **Lwr CI** |
| --- | --- | --- | --- | --- | --- |
|  |  | **GAM_smooth** | 1912.72 | 1922.91 | 1902.53 |
|  |  | **GAM_tensor** | 1912.84 | 1923.67 | 1902.01 |
|  |  | **Bayes** | 1954.67 | 1967.07 | 1942.26 |
|  | **Lifespan** | **LM** | 2069.88 | 2081.61 | 2058.14 |
|  |  | **KNN** | 3126.88 | 3145.94 | 3107.83 |
|  |  | **SVM** | 4800.91 | 4824.54 | 4777.28 |
|  |  | **RF** | 5978.74 | 6024.63 | 5932.85 |
| **Peak** |  | **GBoost** | 10919.99 | 11148.43 | 10691.55 |
| **Area (intakes)** |  | **LM** | 1932.41 | 1944.72 | 1920.10 |
|  |  | **GAM_smooth** | 2022.98 | 2037.21 | 2008.76 |
|  |  | **GAM_tensor** | 2032.59 | 2046.05 | 2019.13 |
|  | **Reproductive** | **Bayes** | 2041.78 | 2055.65 | 2027.91 |
|  | **rate** | **KNN** | 6181.58 | 6216.59 | 6146.57 |
|  |  | **RF** | 11001.14 | 11158.15 | 10844.12 |
|  |  | **SVM** | 14614.46 | 14674.16 | 14554.76 |
|  |  | **GBoost** | 16358.27 | 16555.54 | 16161.00 |
|  |  | **LM** | 2201.23 | 2213.00 | 2189.47 |
|  |  | **GAM_tensor** | 2208.55 | 2220.72 | 2196.38 |
|  |  | **Bayes** | 2292.38 | 2310.97 | 2273.80 |
|  | **Lifespan** | **KNN** | 2505.34 | 2570.34 | 2440.33 |
|  |  | **RF** | 4909.45 | 4954.50 | 4864.39 |
|  |  | **SVM** | 5848.48 | 5888.34 | 5808.61 |
|  |  | **GAM_smooth** | 6532.01 | 6587.10 | 6476.93 |
| **Peak** |  | **GBoost** | 9156.89 | 9302.30 | 9011.47 |
| **Area (fixed)** |  | **GAM_smooth** | 2086.55 | 2098.31 | 2074.78 |
|  |  | **SVM** | 2101.93 | 2113.36 | 2090.50 |
|  |  | **GAM_tensor** | 2102.28 | 2112.33 | 2092.24 |
|  | **Reproductive** | **GBoost** | 2145.32 | 2156.60 | 2134.04 |
|  | **rate** | **LM** | 2162.05 | 2171.65 | 2152.45 |
|  |  | **Bayes** | 2211.30 | 2224.25 | 2198.35 |
|  |  | **RF** | 2994.72 | 3025.90 | 2963.55 |
|  |  | **KNN** | 4357.13 | 4377.04 | 4337.22 |

**Table S2. Nutrient spread of the predicted peak region for all models.** All values are given in units of nutrient intake or diet composition (for fixed ratios).

| **Data** | **Trait** | **Model** | **Protein (median)** | **Protein (std)** | **Carb (median)** | **Carb (std)** |
| --- | --- | --- | --- | --- | --- | --- |
|  |  | **LM** | 0.5333 | 0.0858 | 0.1461 | 0.0194 |
|  |  | **Bayes** | 0.5354 | 0.0963 | 0.1446 | 0.0155 |
|  |  | **GAM_tensor** | 0.5576 | 0.0772 | 0.1504 | 0.0218 |
|  | **Lifespan** | **GAM_smooth** | 0.5598 | 0.0752 | 0.1566 | 0.0208 |
|  |  | **RF** | 0.5884 | 0.1184 | 0.2530 | 0.0342 |
|  |  | **KNN** | 0.6239 | 0.0863 | 0.1920 | 0.0253 |
|  |  | **SVM** | 0.6629 | 0.1153 | 0.2202 | 0.0245 |
| **Nutrient** |  | **GBoost** | 0.6721 | 0.1259 | 0.2723 | 0.0339 |
| **spread (intakes)** |  | **LM** | 0.1348 | 0.0214 | 0.0513 | 0.0088 |
|  |  | **Bayes** | 0.1360 | 0.0207 | 0.0512 | 0.0083 |
|  |  | **GBoost** | 0.2006 | 0.0337 | 0.1893 | 0.0412 |
|  | **Reproductive** | **GAM_smooth** | 0.2152 | 0.0260 | 0.0303 | 0.0049 |
|  | **rate** | **GAM_tensor** | 0.2172 | 0.0257 | 0.0290 | 0.0051 |
|  |  | **RF** | 0.2255 | 0.0379 | 0.1280 | 0.0317 |
|  |  | **KNN** | 0.2768 | 0.0287 | 0.1548 | 0.0140 |
|  |  | **SVM** | 0.3541 | 0.0342 | 0.2288 | 0.0240 |
|  |  | **GAM_tensor** | 0.4752 | 0.0980 | 0.0731 | 0.0162 |
|  |  | **RF** | 0.5004 | 0.2436 | 0.0812 | 0.0261 |
|  |  | **KNN** | 0.5376 | 0.0487 | 0.1072 | 0.0505 |
|  | **Lifespan** | **SVM** | 0.6408 | 0.2370 | 0.0795 | 0.0461 |
|  |  | **Bayes** | 0.6416 | 0.1078 | 0.0866 | 0.0146 |
|  |  | **LM** | 0.6502 | 0.1075 | 0.0910 | 0.0159 |
|  |  | **GBoost** | 0.6599 | 0.2394 | 0.1313 | 0.0454 |
| **Nutrient** |  | **GAM_smooth** | 0.8015 | 0.1148 | 0.1080 | 0.0595 |
| **spread (fixed)** |  | **GBoost** | 0.0807 | 0.0662 | 0.0383 | 0.0143 |
|  |  | **GAM_tensor** | 0.1413 | 0.0554 | 0.0229 | 0.0088 |
|  |  | **KNN** | 0.1538 | 0.1273 | 0.0202 | 0.0214 |
|  | **Reproductive** | **Bayes** | 0.1610 | 0.0897 | 0.0228 | 0.0128 |
|  | **rate** | **LM** | 0.1620 | 0.0902 | 0.0206 | 0.0113 |
|  |  | **RF** | 0.2238 | 0.0768 | 0.0231 | 0.0122 |
|  |  | **SVM** | 0.2804 | 0.0765 | 0.0333 | 0.0106 |
|  |  | **GAM_smooth** | 0.4162 | 0.0293 | 0.0444 | 0.0048 |

**Table S3. Nutrigonometry quantification of nutritional trade-offs in developmental time between two developmental temperatures.** Estimates of $\theta_{i,j}$ (in degrees) and $h_{i,j}$ (in g/L). Analysis from the data presented in Kutz et al., 2019. Confidence intervals overlapping zero implies no difference in the peaks. Magnitude of the estimates indicate the strength of nutritional trade-offs (i.e., larger magnitudes indicate stronger nutritional trade-offs). Note that $\theta_{i,j}$ is bound between 0 and 90 degrees (i.e., 0 and $\frac{\pi}{2}$).

| **Parameter** | **Model** | **Mean** | **Sd** | **lwrCI** | **uprCI** |
| --- | --- | --- | --- | --- | --- |
|  | LM | -0.0228 | 2.1983 | -4.332 | 4.286 |
|  | Bayes | -0.0707 | 2.2355 | -4.452 | 4.311 |
| $\theta_{i,j}$ | RF | 0.74892 | 3.1543 | -5.434 | 6.932 |
|  | Gboost | -11.541 | 15.598 | -42.11 | 19.03 |
|  | SVM | -0.0604 | 30.284 | -59.42 | 59.30 |
|  | KNN | -0.0084 | 2.2102 | -4.341 | 4.324 |
|  | LM | 2.0297 | 51.035 | -98.00 | 102.06 |
|  | Bayes | 0.3822 | 50.295 | -98.20 | 98.971 |
| $h_{i,j}$ | RF | 17.283 | 125.803 | -229.31 | 263.88 |
|  | Gboost | 102.95 | 179.903 | -249.68 | 455.60 |
|  | SVM | 0.6422 | 124.636 | -243.67 | 244.95 |
|  | KNN | 5.8109 | 175.197 | -337.61 | 349.23 |
